# The phospholamban pentamer interacts with the sarcoplasmic reticulum calcium pump SERCA

**DOI:** 10.1101/370387

**Authors:** J. P. Glaves, J. O. Primeau, L. M. Espinoza-Fonseca, M. J. Lemieux, H. S. Young

## Abstract

The interaction of phospholamban with the sarcoplasmic reticulum calcium pump (SERCA) is a major regulatory axis in cardiac muscle contractility. The prevailing model involves reversible inhibition of SERCA by monomeric phospholamban and storage of phospholamban as an inactive pentamer. However, this paradigm has been challenged by studies demonstrating that phospholamban remains associated with SERCA and that the phospholamban pentamer is required for cardiac contractility. We have previously used two-dimensional crystallization and electron microscopy to study the interaction between SERCA and phospholamban. To further understand this interaction, we compared small helical crystals and large two-dimensional crystals of SERCA in the absence and presence of phospholamban. In both crystal forms, SERCA molecules are organized into identical anti-parallel dimer ribbons. The dimer ribbons pack together with distinct crystal contacts in the helical versus large two-dimensional crystals, which allow phospholamban differential access to potential sites of interaction with SERCA. Nonetheless, we show that a phospholamban oligomer interacts with SERCA in a similar manner in both crystal forms. In the two-dimensional crystals, a phospholamban pentamer interacts with transmembrane segments M3 of SERCA and participates in a crystal contact that bridges neighboring SERCA dimer ribbons. In the helical crystals, an oligomeric form of phospholamban also interacts with M3 of SERCA, though the phospholamban oligomer straddles a SERCA-SERCA crystal contact. We conclude that the pentameric form of phospholamban interacts with SERCA, and that it plays distinct structural and functional roles in SERCA regulation.

## INTRODUCTION

Membrane transport by the sarco-endoplasmic reticulum Ca^2+^-ATPase (a.k.a. SERCA) is central to restoring cytosolic calcium levels following intracellular calcium release. In muscle, calcium release from the sarcoplasmic reticulum (SR) leads to contraction. During the subsequent relaxation phase, SERCA transports two cytoplasmic calcium ions into the SR in exchange for two to three luminal protons and at the expense of one molecule of ATP. We currently have an excellent understanding of the SERCA transport cycle from crystallographic studies (reviewed in (1-3)). SERCA consists of three cytoplasmic domains (nucleotide-binding, phosphorylation, and actuator domains) connected to a ten-helix transmembrane domain. ATP-binding and phosphoryl-transfer are mediated by the cytoplasmic domains, and the associated conformational changes are coupled to changes in the calcium binding sites located in the transmembrane domain.

Physiologically, SERCA function depends on the differential expression of SERCA isoforms and splice variants, as well as the differential co-expression of small regulatory transmembrane proteins. The SERCA2a isoform is predominantly expressed in cardiac muscle and is regulated by the co-expression of phospholamban (PLN), a 52 amino-acid integral membrane protein that binds to and lowers the apparent calcium affinity of SERCA. In fast-twitch skeletal muscle, SERCA1a is predominantly expressed and is regulated by the co-expression of sarcolipin (SLN), a 31 amino-acid integral membrane protein homologous to PLN. The prevailing model for PLN inhibition of SERCA involves the reversible association of a PLN monomer (4), which binds to the calcium-free state of SERCA and dissociates under conditions of elevated cytosolic calcium or PLN phosphorylation. This model originated with the observation that monomeric forms of PLN – mutants that disrupt the pentameric assembly of PLN – were better inhibitors of SERCA (5,6). Later, cross-linking (7) and co-immunoprecipitation (8) studies suggested that relief of SERCA inhibition was accompanied by the dissociation of PLN. Monomeric and pentameric forms of PLN were said to be in dynamic equilibrium in the membrane, with the pool of monomers and pentamers influenced by the phosphorylation status of PLN (9). Finally, the PLN pentamer was described as an inactive storage form (4,10), despite the absence of direct experimental evidence.

This paradigm for SERCA inhibition by PLN has been challenged by discrepant results, as well as recent studies that show a persistent association between SERCA and PLN. Initial indications were from co-immunoprecipitation studies (8) and EPR spectroscopy (11), which demonstrated that phosphorylated PLN did not dissociate from SERCA as expected. Since then, many studies using FRET (12-14), EPR (15,16), and NMR (17) have reinforced the concept that PLN appears to be a constitutive regulatory subunit of SERCA. In addition, the PLN pentamer is required for normal cardiac development and function (18), despite being described as an inactive storage form. The active roles proposed for the PLN pentamer include a cation selective channel (19,20), modulation of PLN phosphorylation (21,22), and direct interaction with SERCA (23,24). With regard to this latter point, it is interesting to note that the crystal structure of the SERCA-PLN complex contains two PLN molecules in complex with a single SERCA molecule (25). While only one PLN molecule directly interacts with SERCA, the structure clearly reveals that an oligomeric form of PLN interacts with SERCA. Recall that PLN is proposed to exist as a dynamic equilibrium of oligomeric states ranging from pentamer to monomer (9). The crystal structures of the SERCA-PLN and SERCA-SLN complexes revealed a novel calcium-free E1-like state of SERCA (25-27). In these structures, the binding groove for PLN formed by transmembrane helices M2, M6 and M9 of SERCA is shallow enough to accommodate oligomeric forms of PLN (27). Thus, the interaction potential of PLN oligomers with SERCA remains an outstanding question in the field.

Toward this goal, our laboratory previously showed that PLN pentamers interacted with SERCA in two-dimensional (2D) co-crystals in a manner dependent on the functional state of PLN (23,24). The physical interaction in these crystals occurred at an accessory site on SERCA (transmembrane segment M3), yet it bore all the hallmarks normally associated with SERCA inhibition by the PLN monomer – crystal formation depended on the functional, oligomeric and phosphorylation states of PLN. While this suggested that PLN pentamers contribute to SERCA regulation in some unappreciated fashion, uncertainty remained because PLN participates in a crystal contact with SERCA in the large 2D co-crystals. To further examine this SERCA-PLN interaction, we undertook a comparison with helical crystals of the SERCA-PLN complex by electron cryo-microscopy. The site of interaction between SERCA and PLN in the large 2D crystals – the M3 accessory site of SERCA – is involved in a SERCA-SERCA crystal contact in the helical crystals. Since the SERCA-SERCA contact facilitates helical crystal formation, a PLN interaction at this site is not required for crystal formation. Nonetheless, we found transmembrane densities in the helical crystals that were attributable to PLN and associated with M3 of SERCA. We also demonstrated that the functional effects of PLN on SERCA depend on the stoichiometry, suggesting that PLN oligomers have a unique functional effect on SERCA. Combined, the results support an interaction between PLN pentamers and SERCA, with functional consequences that are distinct from the inhibitory interaction. A molecular model of this SERCA-PLN pentamer complex was generated using protein docking and molecular dynamics simulations.

## MATERIALS & METHODS

### Materials

All reagents were of the highest purity available: octaethylene glycol monododecyl ether (C_12_E_8_; Barnet Products, Englewood Cliff, NJ); egg yolk phosphatidylcholine (EYPC), phosphatidylethanolamine (EYPE) and phosphatidic acid (EYPA) (Avanti Polar Lipids, Alabaster, AL); all reagents used in the coupled enzyme assay including NADH, ATP, PEP, lactate dehydrogenase, and pyruvate kinase (Sigma-Aldrich, Oakville, ON Canada).

### Co-reconstitution of PLN and SERCA

Recombinant PLN was expressed and purified as previously described (28). Lyophilized PLN (100 μg) was suspended in a 100 μl mixture of chloroform-trifluroethanol (2:1) and mixed with lipids (400 μg EYPC, 50 μg EYPE, 50 μg EYPA) from stock chloroform solutions. The peptide-lipid mixture was dried to a thin film under nitrogen gas and desiccated under vacuum overnight. The peptide-lipid mixture was hydrated in buffer (20 mM imidazole pH 7.0; 100 mM KCl; 0.02% NaN_3_) at 37 °C for 10 min, cooled to room temperature, and detergent-solubilised by the addition of C_12_E_8_ (0.2 % final concentration) with vigorous vortexing. Detergent-solubilized SERCA was added (500 μg in a total volume of 500 μl) and the reconstitution was stirred gently at room temperature. Detergent was slowly removed by the addition of SM-2 biobeads (Bio-Rad, Hercules, CA) over a 4-hour time course (final weight ratio of 25 biobeads to 1 detergent). Following detergent removal, the reconstitution was centrifuged over a sucrose-gradient for 1 h at 100,000*g*. The resultant layer of reconstituted proteoliposomes was removed, flash-frozen in liquid-nitrogen and stored at −80 °C. The final approximate molar ratios were 120 lipids to 3.5 PLN to 1 SERCA.

### Crystallization conditions

Co-reconstituted proteoliposomes were collected by centrifugation in crystallization buffer (20 mM imidazole, pH 7.4, 100 mM KCl, 5 or 35 mM MgCl_2_, 0.5 mM EGTA, 0.25 mM Na_3_VO_4_, with or without 30 μM thapsigargin). The pellet was subjected to two freeze-thaw cycles, resuspended with a micropipette, followed by two additional freeze-thaw cycles. The samples were incubated at 4°C for several days to one week. The use of 5 mM MgCl_2_ in the crystallization buffer promoted the formation of helical crystals, while the use of 35 mM MgCl_2_ promoted the formation of large 2D crystals. The large 2D crystals were formed from SERCA in the presence of a Lys^27^-Ala PLN mutant (human PLN sequence (24)), while the helical crystals were formed from SERCA in the presence of an Asn^27^-Ala PLN mutant (canine PLN sequence (29)). Note that these two sequences are nearly identical with two minor sequence variations (Glu^2^ and Lys^27^ in human PLN are Asp^2^ and Asn^27^ in canine PLN). Thus, the PLN variants in the crystals differ only by having Glu^2^ or Asp^2^, and they are both super-inhibitory, pentameric forms of PLN.

### ATPase activity assays of SERCA reconstitutions

ATPase activity of the co-reconstituted proteoliposomes was measured by a coupled-enzyme assay over a range of calcium concentrations from 0.1 μM to 10 μM (30,31). The V_max_ (maximal velocity), K_Ca_ (apparent calcium affinity) and n_H_ (Hill coefficient) were determined by fitting the data to the Hill equation (Sigma Plot software, SPSS Inc., Chicago, IL). Errors were calculated as the standard error of the mean for a minimum of four independent reconstitutions.

### Data processing

The helical crystals datasets have been published (**Table 1** & (29,32)). The structure from helical crystals of SERCA alone was an average generated from five helical symmetry groups (−22, 6 reference symmetry). The structure from helical crystals of SERCA in the presence of PLN (Asn^27^-Ala mutant; canine sequence) was an average generated from two helical symmetry groups (−27, 9 reference symmetry). Density maps were calculated for the SERCA-PLN helical crystals without enforcing two-fold symmetry. The two molecules composing the unit cell for the control structure (SERCA alone) were masked and aligned by cross-correlation with the corresponding molecules from the SERCA-PLN map. After alignment, the two molecules composing the unit cell for the control structure were summed to generate a single map and back-transformed to Fourier space to generate layer-line data with the SERCA-PLN helical symmetry (−27, 9 reference symmetry). Density maps were then calculated from the control and PLN layer line data with and without two-fold symmetry enforced. Since the control and PLN layer line data were now of the same helical symmetry group, a difference map could be calculated by simple density scaling and pixel-by-pixel subtraction.

**Table 1:** Crystallographic data for frozen-hydrated helical crystals of SERCA in the absence and presence of PLN is summarized.

|  | SERCA<sup>a</sup> | SERCA-PLN<sup>b</sup> |
| --- | --- | --- |
| Number of crystals | 58 | 32 |
| Helical symmetries | 5 | 2 |
| Reference symmetry | -22, 6 | -27, 9 |
| Twofold phase statistics<sup>c</sup> |  |  |
| 40 Å | 1.1° | 0.7° |
| 30 Å | 1.7° | 1.7° |
| 20 Å | 4.0° | 3.8° |
| 14 Å | 15.2° | 12.5° |
| 10 Å | 27.5° | 23.8° |
| 8 Å | 37.2° | 38.8° |

<sup>a</sup>Previously described by Xu et al (2002)
<sup>b</sup>Previously described by Young et al (2001)
<sup>c</sup>Amplitude-weighted phase residuals for twofold symmetry including 100% of the off-equatorial data for the indicated resolution ranges. Random phases produce a phase residual of 45°.

### Atomistic modeling of the PLN pentamer bound to SERCA

The crystal structure of SERCA1a in the E2·MgF_4_^2-^ state (PDB accession code 1WPG) and the NMR structure of the PLN pentamer (PDB accession code 2KYV) were used as templates for the construction of an atomic model of the SERCA-PLN complex. Protein-protein docking simulations identified the binding interface between the M3 helix of SERCA and the PLN pentamer. The program ClusPro (33) was used to dock the M3 helix (residues Pro^248^ to Phe^279^ were included) to the transmembrane helices of the PLN pentamer (residues Gln^26^ to Leu^52^ were included). Only residues that do not participate in intramolecular SERCA contacts were considered as potential SERCA-PLN interaction sites (i.e. Glu^258^, Gln^259^, Lys^262^, Leu^266^, Val^269^, Trp^272^, Leu^273^, Ile^276^, and Phe^279^). The resulting docked orientations were clustered and compared against the 2D crystallographic data to select the most appropriate M3-PLN pentamer complex. The SERCA-PLN pentamer complex was then constructed by superposing the coordinates of the M3 helix onto the crystal structure of the E2·MgF_4_^2-^ state of SERCA. The final model was subjected to 5,000 steps of energy minimization.

### Molecular dynamics simulations of the SERCA-PLN pentamer complex

The atomistic SERCA-PLN pentamer complex served as a starting structure to obtain a structural model of the complex at physiologically relevant simulation conditions. Based on previous studies of the E2 state of SERCA (34), we modeled transport site residues Glu^309^, Glu^771^ and Glu^908^ as protonated and residue Asp^800^ as ionized. The complex was inserted in a pre-equilibrated 120 Å^2^ bilayer of POPC lipids. This lipid-protein system was solvated using TIP3P water molecules with a minimum margin of 15 Å between the protein and the edges of the periodic box in the *z*-axis, and K^+^ and Cl^-^ ions were added (KCl concentration of ~0.1 mM).

To generate a reliable model of the complex, we performed molecular dynamics (MD) simulations of the fully solvated complex using NAMD 2.12 (35) with periodic boundary conditions (36), particle mesh Ewald (37,38), a non-bonded cut-off of 9 Å, and a 2 fs time step. The CHARMM36 force field topologies and parameters were used for the proteins (39), lipid (40), water, and ions. The NPT ensemble was maintained with a Langevin thermostat (310K) and an anisotropic Langevin piston barostat (1 atm). The system was first subjected to energy minimization, followed by gradually warming up of the system for 200 ps. This procedure was followed by 10 ns of equilibration with backbone atoms harmonically restrained using a force constant of 10 kcal mol^-1^ Å^-2^. The MD simulation of the complex was continued without restraints for 200 ns.

## RESULTS

### Functional differences between the PLN monomer and pentamer

The PLN-SERCA molar ratio reported for cardiac SR membranes varies from 1:1 (41) to 4:1 (42-44), suggesting that modulation of PLN content in the SR membranes may play a role in the regulation of SERCA and cardiac contractility. Yet the effect of PLN on the apparent calcium affinity of SERCA saturates at a molar ratio of approximately one PLN monomer per SERCA (42,45). This raises the question what are the consequences of excess PLN in cardiac SR membranes? To investigate this, we measured the calcium-dependent ATPase activity of SERCA alone and in the presence of two molar ratios of SERCA-PLN (1:2.5 and 1:5; **Figure 1**). These SERCA-PLN ratios were chosen to saturate the effect of the PLN monomer (1:2.5 ratio) and reveal any effects of the PLN pentamer at a higher ratio (1:5). While the effect of PLN on the apparent calcium affinity (K_Ca_) of SERCA remained unchanged under these conditions, there was a statistically significant increase in the maximal activity (V_max_) of SERCA at the 1:5 molar ratio of SERCA-PLN. The V_max_ values for SERCA alone (4.02 ± 0.04 μmoles/min/mg), SERCA with 2.5 moles of PLN (5.25 ± 0.13 μmoles/min/mg) and SERCA with 5 moles of PLN (6.03 ± 0.08 μmoles/min/mg) were all statistically significant from one another (p<0.01). This result supports the notion that SERCA activity is influenced by the membrane concentration of PLN (31,46,47). The dependence of V_max_, but not K_Ca_, on the SERCA-PLN membrane concentration suggests that there are multiple modes of interaction between PLN and SERCA, involving more than a simple one-to-one inhibitory interaction between these proteins.

**Figure 1:**
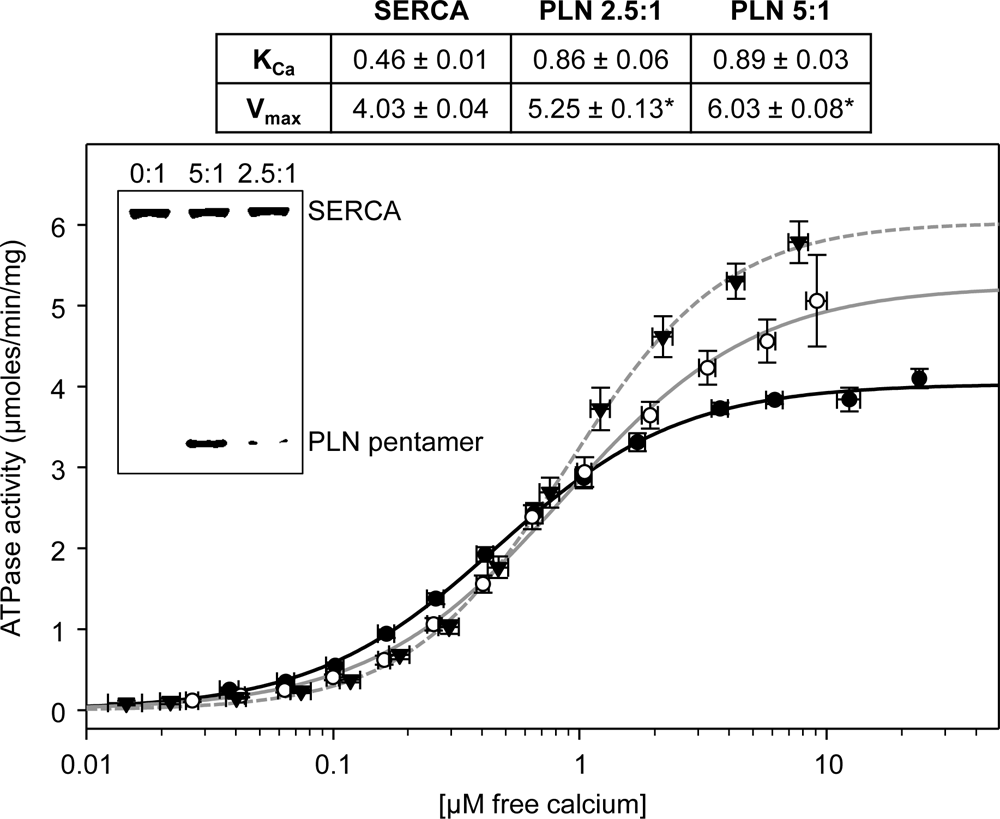
ATPase activity as a function of calcium concentration for SERCA proteoliposomes in the absence and presence of PLN. Proteoliposomes containing SERCA alone (filled circles, black line), proteoliposomes containing SERCA and a 2.5-molar ratio of PLN (open circles, grey line), and proteoliposomes containing SERCA and a 5-molar ratio of PLN (filled triangles, grey dashed line). The lines represent curve fitting of the experimental data using the Hill equation. The kinetic parameters are indicated in the inset **Table**, and the asterisk (*) indicates statistical significance (p<0.01) compared to SERCA alone. The V_max_ values in the presence of PLN are also statistically significant from one another (p<0.01). Each data point is the mean ± SEM (*n* ≥ 4). ***Inset –*** 10% SDS-PAGE gel of the reconstituted proteoliposomes of SERCA in the absence and presence of PLN. The bands corresponding to SERCA and the PLN pentamer are indicated. The molar ratio of PLN-to-SERCA is indicated for each lane (0:1, 5:1, and 2.5:1).

### Crystallization

Our previous studies of large 2D crystals of the SERCA-PLN complex identified an M3 accessory site with characteristics consistent with a functional interaction (23,24). However, these observations contradicted the central dogma that the pentamer is an inactive storage form of PLN. Because PLN also participated in a crystal contact in the 2D crystals, the relevance of this interaction remained ambiguous. Thus, it became important to determine if the interaction between SERCA and the PLN pentamer is a crystal contact or a natural association. To address this issue, we re-examined helical crystals of the SERCA-PLN complex (29,32,48) and compared them to the large 2D crystals (23,24) (**Figure 2**). Helical crystals and large 2D crystals are grown from reconstituted proteoliposomes containing either SERCA alone or SERCA in the presence of PLN. The proteoliposomes readily form crystals when incubated in decavanadate and EGTA buffer (20 mM imidazole, pH 7.4, 100 mM KCl, 5 or 35 mM MgCl_2_, 0.5 mM EGTA, and 0.25 mM Na_3_VO_4_), followed by several freeze-thaw cycles to promote proteoliposome fusion and the growth of large crystals. Incubation at 4°C produces optimal crystal formation within three to five days. Importantly, these experimental conditions are identical for the formation of helical and large 2D crystals except for the concentration of magnesium – 5 mM MgCl_2_ promoted the formation of helical crystals and 35 mM MgCl_2_ promoted the formation of large 2D crystals. The fundamental units of both crystal forms are the well characterized anti-parallel dimer ribbons of SERCA molecules (23,32), which are known to be a stable structural element induced by decavanadate in the absence of calcium (49). However, the SERCA dimer ribbons are organized differently in the helical and large 2D crystals (**Figure 3**). Important for this study, the organization of the SERCA dimer ribbons in the helical crystals had a large impact on the pentamer interaction site observed in the large 2D crystals.

**Figure 2:**
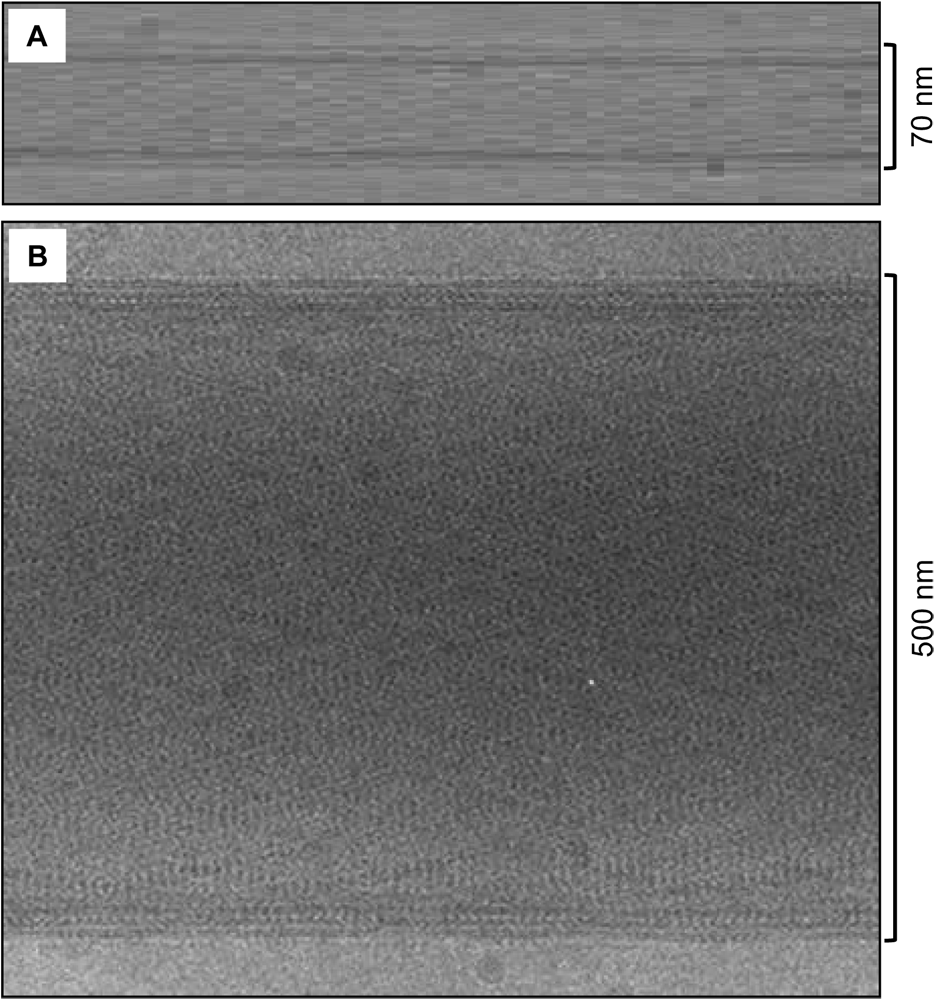
Representative examples of frozen-hydrated helical crystals (A) and large 2D crystals (B) of SERCA and PLN.

**Figure 3:**
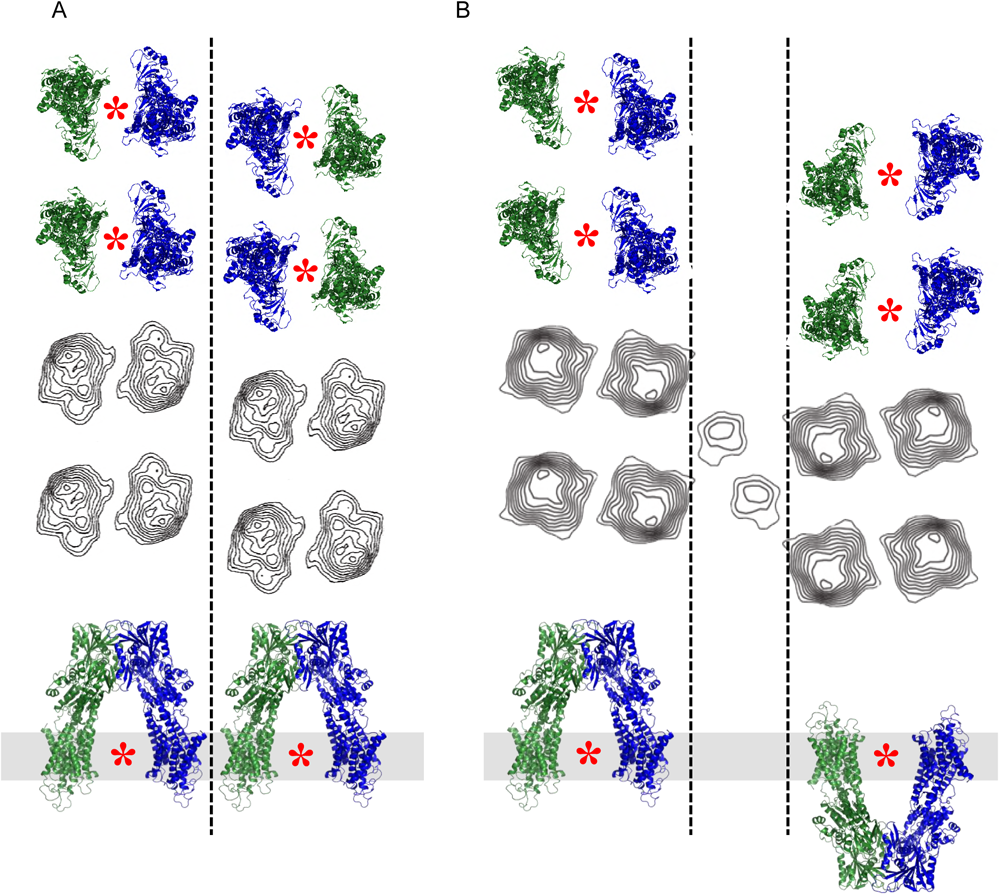
Pseudo-atomic models and projection maps for the helical crystals (A) and large 2D crystals (B) of SERCA and PLN. The SERCA molecules are arranged in anti-parallel dimer ribbons in both lattices. ***Top panels*** are the view orthogonal the membrane bilayer (PDB accession code 1IWO); ***Bottom panels*** are the view normal to the membrane bilayer. The asterisks indicate the location of the inhibitory binding site for PLN (M2, M6 & M9 of SERCA), which is open in both crystal forms. The dashed line indicates the location of the accessory site for PLN binding (M3 of SERCA), which is limited in the helical crystals and open in the large 2D crystals. ***Middle panels*** are the projection contour maps for helical crystals (A) and large 2D crystals (B). The additional densities in the projection map of the large 2D crystals correspond to PLN pentamers (between the dashed lines; the density for two pentamers is shown).

### Helical versus large 2D crystals

In both crystal forms, the native-like membrane environment is well suited for structural studies of SERCA regulatory complexes (23,24,29). The helical crystals are membrane cylinders with a relatively narrow diameter (~70 nm), where the SERCA dimer ribbons arranged with p2 symmetry (**Figure 3**). All SERCA molecules are oriented with their large cytoplasmic domains on the exterior of the membrane cylinder, and there is no gap between neighboring SERCA dimer ribbons. The important point here is that the inhibitory binding site for PLN (M2, M6 & M9 of SERCA) is unobstructed, while the accessory site (M3) is involved in a SERCA-SERCA crystal contact that brings together neighboring dimer ribbons in the helical lattice (**Figure 3**).

The large 2D crystals form as single or double-layered membrane tubes with a relatively large diameter (0.5 to 3 μm). The 2D crystals are composed of the same SERCA dimer ribbons that make up the helical crystals, yet they are arranged differently in a p22_1_2_1_ lattice (**Figure 3** (23)). The SERCA molecules are oriented with their cytoplasmic domains alternately extending from both sides of the membrane, and there is a large gap between neighboring SERCA dimer ribbons. Thus, both the inhibitory binding site for PLN (M2, M6 & M9 of SERCA) and the accessory site (M3) are open and unobstructed (**Figure 3**). In comparing the two crystals forms, the inhibitory site on SERCA is accessible in both helical and large 2D crystals, while a SERCA-SERCA crystal contact limits access to the M3 accessory site in the helical crystals.

### Helical crystals

In the present study, we reanalyzed the helical crystals of SERCA and PLN (29) in order to identify densities associated with the transmembrane domain of PLN. Recall that our prior studies of the regulatory SERCA-PLN complex using helical crystals at 10 Å resolution provided a well-defined three-dimensional structure for SERCA and weak, discontinuous densities attributable to the cytoplasmic domain of PLN (29). To observe these PLN densities, it was necessary to calculate difference maps between helical reconstructions of SERCA in the absence and presence of PLN. Nonetheless, a model was proposed in which the cytoplasmic domain of PLN interacted with two SERCA molecules, though no difference densities were observed for the transmembrane helix of PLN (29).

To improve upon our previous analyses, we hypothesized that a higher-resolution control would facilitate comparison with our SERCA-PLN datasets and lead to the identification of transmembrane densities. A 6 Å resolution structure of SERCA by electron cryo-microscopy was reported, which improved upon our previous control dataset by increasing the number of helical crystals included in the analysis (**Table 1** & (32)). The quality of the resultant density map was sufficient for fitting of SERCA atomic coordinates, which required rigid-body docking of SERCA domains into corresponding regions of the map. This higher-resolution control dataset (SERCA in the absence of PLN) was necessary for the visualization of PLN densities. Second, to identify transmembrane densities, we focused on the highest resolution dataset for SERCA in the presence of PLN (29). This high-resolution data set of SERCA-PLN helical crystals was enabled by (i) the use of a super-inhibitory pentameric form of PLN (Asn^27^-Ala mutant; canine PLN sequence), (ii) a one-to-3.5 molar ratio of SERCA to PLN, and (iii) the absence of thapsigargin, which has traditionally been used to promote helical crystals.

To determine structural differences in the presence of PLN, we focused on helical crystals of the super-inhibitory pentameric form of PLN (Asn^27^-Ala mutant). The crystallographic data for the control and Asn^27^-Ala datasets have similar two-fold phase statistics as a function of resolution to 8 Å (**Table 1**). The helical reconstructions for SERCA in the absence and presence of PLN at 8 Å resolution were subjected to two rounds of density scaling and real space alignment, followed by density subtraction to produce a difference map (**Figures 4 & 5**). This comparison finally revealed densities attributable to the transmembrane domain of PLN, which could be interpreted in terms of a pseudo-atomic model for SERCA in the helical crystals (32). A set of densities were found at the M3 accessory site, along a twofold axis that relates a pair of SERCA molecules from neighboring dimer ribbons. The twofold axis corresponds to a crystal contact between SERCA molecules involving M3-M4 loops, where four PLN densities traverse the membrane and straddle a pair of M3 segments (**Figure 4**). Similarly, a series of densities were observed at the surface of the membrane, directly above the transmembrane densities (**Figure 5**). While the densities were consistent with a PLN tetramer, the observed oligomeric state is likely influenced by the twofold axis formed by the SERCA-SERCA crystal contact. An additional weak density was observed adjacent to the PLN tetramer, suggesting partial occupancy of a fifth PLN molecule (i.e. a PLN pentamer; **Figure 4**, arrow). Nonetheless, a PLN oligomer was found associating with M3 of SERCA in the helical crystals, which is precisely what was observed in large 2D co-crystals of the SERCA-PLN complex (23,24).

**Figure 4:**
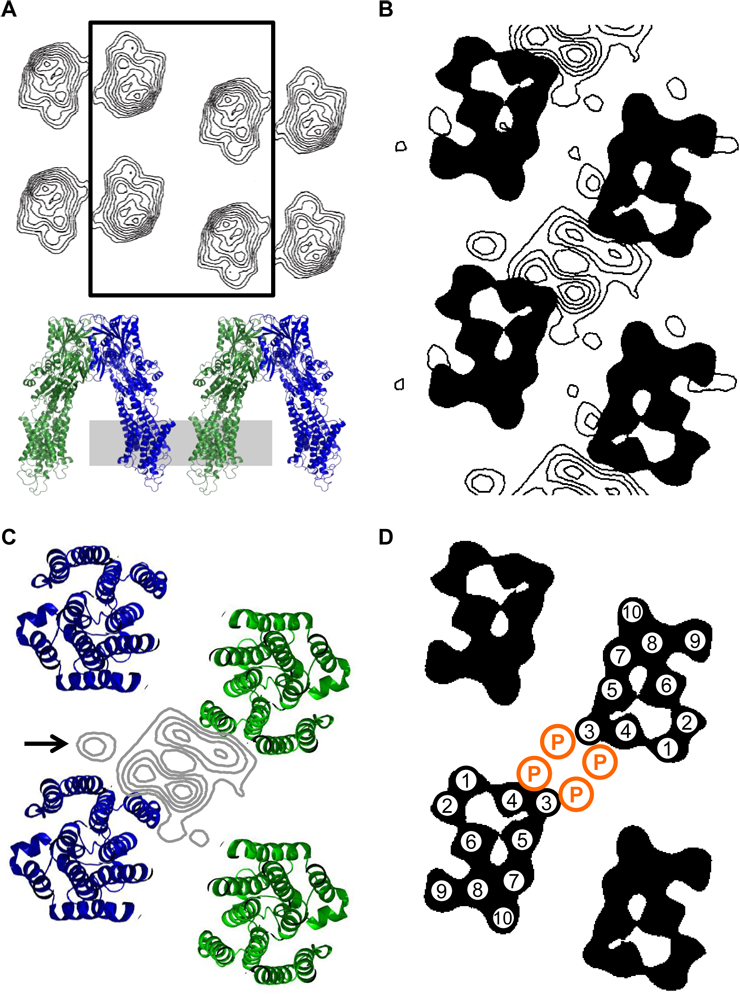
Transmembrane difference density map from helical crystals of SERCA in the absence and presence of PLN. (A) Projection map for the helical crystals with a box indicating the area of the helical lattice shown in the following panels, and a pseudo-atomic model with a grey bar indicating the section of the map shown for the difference densities. (B) Difference density map at 8 Å resolution revealing transmembrane densities attributable to PLN (grey contours; 0.75 σ increment). The organization of SERCA’s ten transmembrane helices are shown as a black silhouette (four SERCA molecules are shown). (C) Pseudo-atomic model of the transmembrane domain, shown for four symmetry-related SERCA molecules in the helical crystals (viewed normal to the membrane bilayer). The difference density map (grey contours) is overlaid on the transmembrane helices of SERCA. Four discrete densities are observed for PLN; there is a weak fifth density adjacent to these densities (arrow). (D) The positions of the ten transmembrane segments of SERCA and the PLN transmembrane helices (orange circles) as indicated by the difference density map shown in panels (B) and (C).

**Figure 5:**
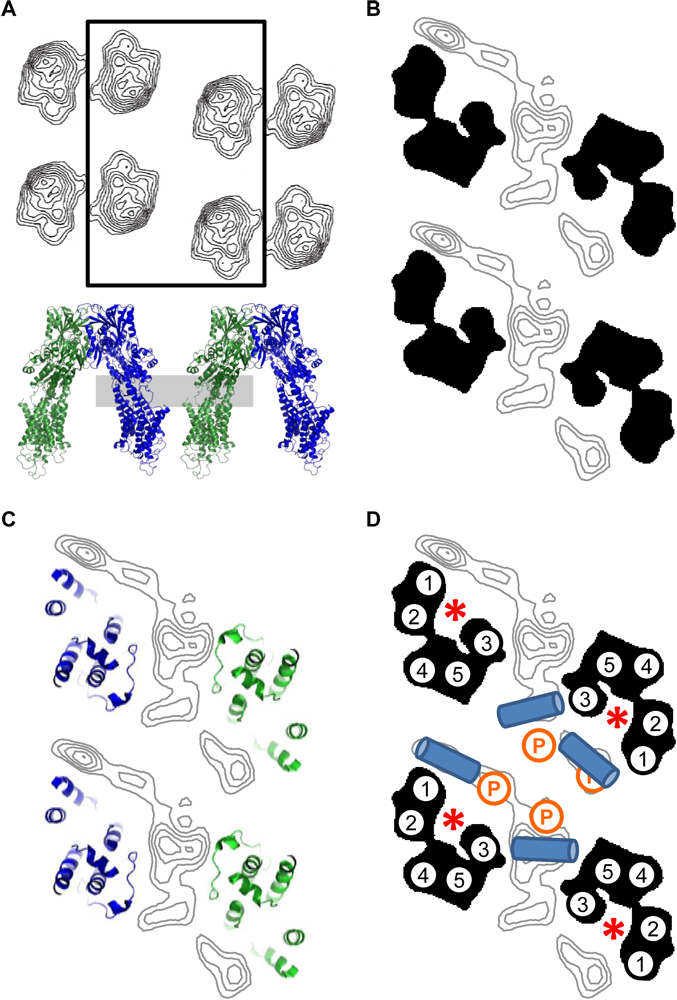
Difference density map at the surface of the membrane from helical crystals of SERCA in the absence and presence of PLN. (A) Shown again for clarity is the projection map for the helical crystals with a box indicating the area of the helical lattice shown in the following panels. The pseudo-atomic model is also shown with a grey bar indicating the section of the map corresponding to the difference densities. (B) Difference density map at 8 Å resolution revealing discrete densities attributable to the cytoplasmic domain of PLN (grey contours; 0.75 σ increment). The organization of SERCA’s “stalk” region is shown as a black silhouette (four SERCA molecules are shown). (C) Pseudo-atomic model for this region of SERCA, showing four molecules in the helical lattice viewed normal to the membrane bilayer. The difference density map (grey contours) is overlaid for comparison. There are two sets of four discrete densities observed for PLN within this region of the map. (D) The positions of the helical segments of SERCA and the PLN cytoplasmic domains (only the central four are shown as blue cylinders) as indicated by the difference density map shown in panels (B) and (C). The positions of the PLN transmembrane densities are also indicated (orange circles).

### Molecular model of the SERCA-PLN pentamer complex

To generate a molecular model of the SERCA-PLN pentamer complex that was representative of the state found in the 2D crystals, we performed protein-protein docking and MD simulations. Only the transmembrane domains of the PLN pentamer were included in the docking and MD simulations. Following the protein-protein docking, the SERCA-PLN complex that most closely matched the arrangement in the 2D crystals was selected. This complex was embedded in a lipid bilayer (1-palmitoyl-2-oleoyl-glycero-3-phosphocholine; POPC), fully hydrated to mimic physiological conditions, and subjected to 200ns MD simulations. This latter step was done in order to validate the stability of the complex and ensure appropriate packing constraints at the molecular interface between SERCA and the PLN pentamer. We found that transmembrane segment M3 of SERCA interacts with the PLN pentamer at the interface between two PLN monomers. The interaction occurs along the entire length of the PLN pentamer and M3 of SERCA (**Figure 6** & Supplemental Figure 1), largely stabilized by hydrophobic interactions (e.g. Ile^45^ of PLN and Trp^272^ of SERCA; not shown).

**Figure 6:**
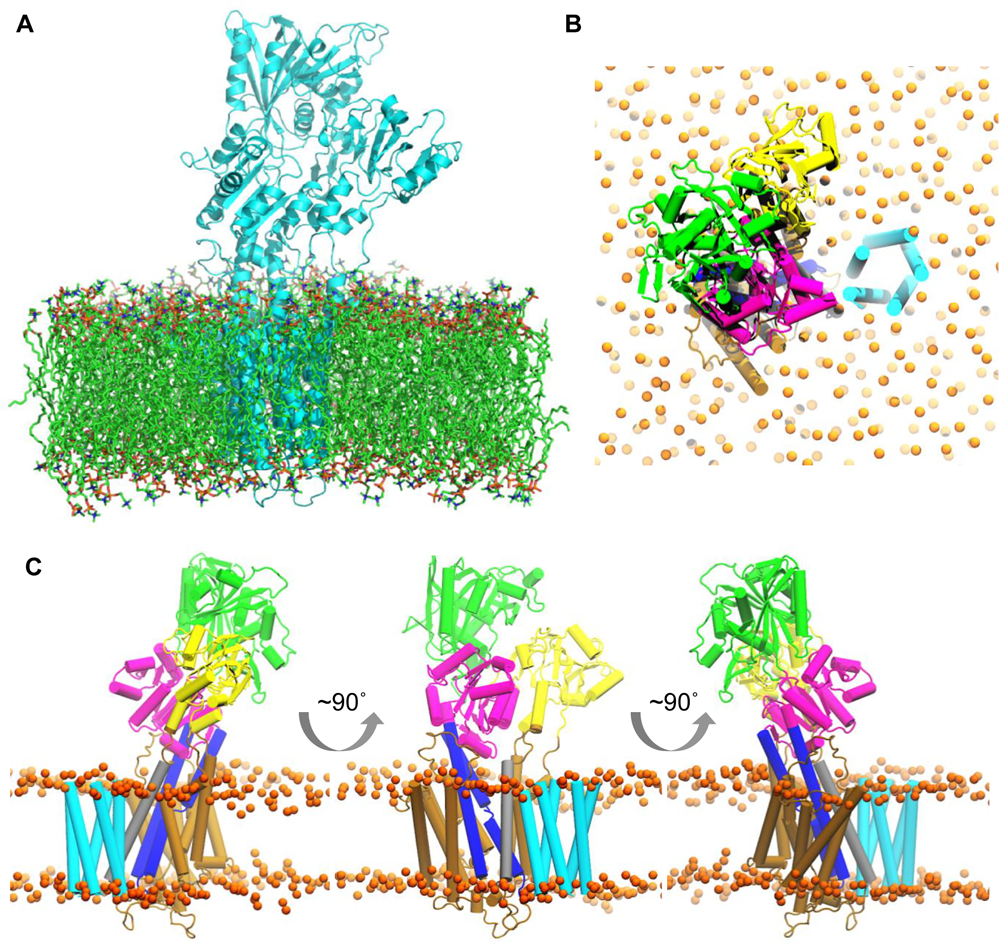
Molecular model for the interaction of SERCA with the pentameric form of PLN. (A) The full atomistic model generated from protein-protein docking and MD simulations is shown. SERCA and PLN are shown in cartoon representation (cyan) and the lipid bilayer is shown in stick representation. (B) Molecular model for the interaction of the PLN pentamer with the M3 accessory site of SERCA, viewed along the membrane surface. The nucleotide binding domain (green), the phosphorylation domain (magenta), the actuator domain (yellow), transmembrane segments M4 and M5 (blue), transmembrane segment M3, and the transmembrane domain of the PLN pentamer are indicated. The lipid headgroups are shown as orange spheres. (C) Three orthogonal views of the complex are shown with the actuator domain of SERCA in the foreground (left), M3 of SERCA in the foreground (middle), and the nucleotide binding and phosphorylation domains in the foreground (right).

## DISCUSSION

Early studies of PLN focused on the monomeric state as the species responsible for SERCA inhibition (5,6), while the pentamer was assigned the default role of an inactive storage form. This was based on the observation that periodic mutations in the transmembrane domain of PLN disrupt the pentamer, enhance monomer formation, and increase SERCA inhibition. The consensus model that emerged included SERCA inhibition by monomeric PLN, a reversible association between PLN and SERCA (50), and the dissociation of PLN as the primary mechanism for relieving SERCA inhibition (i.e. following phosphorylation of PLN or elevated cytosolic calcium). Molecular models of the SERCA-PLN inhibitory complex, based on the calcium-free E2 state of SERCA, also appeared to support the monomer as the inhibitory form of PLN. In the E2 state, there is a deep groove formed by transmembrane segments M2, M6 and M9 of SERCA. Modeling PLN’s transmembrane helix into this groove provided a plausible mechanism for SERCA inhibition (51,52). PLN would impede groove closure and the E2-to-E1 transition that accompanies calcium binding by SERCA. In these molecular models, the deep binding groove could not accommodate a PLN pentamer. However, recent crystal structures of the SERCA-PLN (25) and SERCA-SLN (26,27) complexes have shed new light on the regulatory interaction. The structure of SERCA in these complexes is a novel E1-like, calcium-free state with PLN or SLN bound in a partially closed groove formed by M2, M6 and M9 of SERCA. In the structure of the SERCA-PLN complex, there are two PLN transmembrane helices associated with SERCA, and the shallow binding groove could accommodate a PLN pentamer (27). Thus, the consensus model no longer adequately describes all of the available evidence regarding SERCA-PLN interactions.

### Functional consequences of PLN oligomers

In the present study, the proteoliposomes containing SERCA and PLN provided a well-defined and well-characterized (23,24,29,46,53) starting material for functional analysis and crystal formation. Like cardiac and skeletal muscle SR membranes, the proteoliposomes are densely packed with SERCA molecules uniformly oriented with their cytoplasmic domains on the exterior side of the lipid membrane (molar ratio of ~120 lipids per SERCA (53)). The proteoliposomes also contain a molar excess of PLN (~2.5 to 5 molecules of PLN per SERCA), where the majority of PLN is similarly oriented with the cytoplasmic domains on the exterior side of the membrane (31). These SERCA-PLN ratios are similar to cardiac SR membranes (42-44). PLN exists as both a monomer and a pentamer, with approximately 80% of PLN in the pentameric state (5,9,24,31,54,55). Thus, the stoichiometry in the proteoliposomes ranges from 0.4 to 0.8 PLN pentamers per SERCA, respectively. These proteoliposomes were first used to measure SERCA ATPase activity in the absence and presence of PLN (**Figure 1**). Two molar ratios of PLN were chosen to demonstrate the dependence of SERCA function on increasing concentrations of the PLN pentamer. We observed an increase in the maximal activity of SERCA at the higher concentration of PLN pentamers. In contrast, the effect of PLN on the apparent calcium affinity of SERCA saturated at the lower concentration of PLN pentamers, as expected. Thus, the effect on the maximal activity of SERCA is dependent on the concentration of PLN pentamers and separate from the inhibitory interaction between PLN and SERCA.

### Structural consequences of PLN oligomers

The same proteoliposomes used for functional assays were used to generate helical and large 2D crystals of SERCA in the absence and presence of PLN, which formed under relatively simple crystallization conditions. Besides magnesium discussed above, the essential components included EGTA to remove calcium and decavandate to induce formation of the SERCA dimer ribbons. The removal of calcium promotes an E2, calcium-free state of SERCA that is compatible with PLN binding. Decavanadate interacts with SERCA at two sites, one that encompasses the nucleotide binding site and a second site that bridges the actuator domains of two SERCA molecules and thereby stabilizes the antiparallel dimer ribbons (56).

Our initial finding that PLN pentamers interacted with SERCA in large 2D crystals was a surprise (23). At the time, the simplest explanation was that the PLN pentamer makes a crystal contact with SERCA that is not relevant to the function of these two proteins. However, subsequent studies revealed that the SERCA-PLN interaction in the crystals responded to physiological perturbations (24) – namely, phosphorylation of PLN and mutations that impact PLN function also impeded crystal formation. These perturbations have been used as hallmarks for the functional interaction between SERCA and PLN. Thus, the PLN pentamer appeared to interact with SERCA in a functionally-relevant, though unexpected manner. However, there were two SERCA interaction sites identified in the crystal lattice, one with M3 and one with M10 of SERCA, and the PLN pentamer clearly participated in a crystal contact. This ambiguity led us to consider another crystal form capable of discriminating between a crystallographic and non-crystallographic interaction. The helical crystals of the SERCA-PLN complex satisfied this requirement. A simple change in the crystallization conditions – 5 mM versus 35 mM magnesium– dramatically altered the crystal packing and the relative accessibility of transmembrane segment M3 of SERCA. The helical crystals form at a lower magnesium concentration (5 mM), and they transform into the large 2D crystals at a higher magnesium concentration (35 mM).

With this small change in magnesium, a major difference between the helical and large 2D crystals was the accessibility of transmembrane segment M3 of SERCA. In the helical crystals, M3 is involved in a SERCA-SERCA crystal contact, which is absent in the large 2D crystals. Thus, the helical crystals offer limited access to the M3 accessory site, while this site is accessible in the large 2D crystals. Despite the limited access to M3 in the helical crystals, an oligomeric form of PLN was found straddling the M3 transmembrane segments of adjacent SERCA molecules (**Figure 4**). Four discreet transmembrane densities were found, which correspond to the transmembrane helices of a PLN tetramer. The evidence for the densities corresponding to PLN include: (i) the size and density of each peak (~3.5 σ) are consistent with transmembrane helices; (ii) besides SERCA, the only other protein present in the co-reconstituted proteoliposomes is PLN (at ~1-to-3.5 SERCA-PLN molar ratio); and (iii) we know that PLN is present in the crystal lattice because an anti-PLN monoclonal antibody disrupts crystallization (29). We observed four densities rather than the five expected for a PLN pentamer; however, the PLN oligomer sits across a two-fold axis and this must influence the interaction and the observed densities. There is an adjacent fifth density (**Figure 4**), though it is weak and may represent partial occupancy of an additional PLN molecule. We also observed four densities at the surface of the membrane (equivalent to the cytosolic side of the SR membrane), situated above each of the PLN transmembrane densities (**Figure 5**). These densities are consistent with the N-terminal α-helix of PLN and molecular models that place it at the membrane surface (17,57). In this case, the PLN oligomer does not appear to interact with M10 of SERCA or another PLN oligomer, as was observed in the large 2D crystals. Thus, the data presented herein indicate that PLN associates with the M3 accessory site of SERCA independent of crystal contacts (**Figure 6**).

### The PLN pentamer and SERCA activity

The increased maximal activity of SERCA in the presence of PLN pentamers has been shown to be sensitive to mutation and the oligomeric state of PLN (46,47). Here we show that the increase in the maximal activity of SERCA depends on the abundance of PLN pentamers in the same proteoliposomes used for 2D crystallization (**Figure 1**). The maximal activity of SERCA was measured for each proteoliposome preparation to confirm the functional effect of the PLN pentamer, and the remainder of the proteoliposomes were subjected to crystallization trials. Thus, we can consider the interaction in the helical and large 2D crystals in the context of the effect on the maximal activity of SERCA.

There are two well-known mechanisms for increasing the maximal activity of SERCA. The first is to solubilize SERCA in the detergent C_12_E_8_ (58,59), which frees SERCA from the lateral constraints of the lipid bilayer and increases its maximal activity approximately two-fold. The second is to reconstitute SERCA in the presence of a mixture of lipids including phosphatidylcholine, phosphatidylethanolamine (PE), and phosphatidic acid (PA). The addition of lipids that mildly destabilize the bilayer (PE & PA) also increase SERCA’s maximal activity (53,60). With this context in mind, the observed physical interaction between the PLN pentamer and SERCA offers a plausible explanation for the stimulation of SERCA’s maximal activity. The M3 accessory site lines the calcium entry funnel of SERCA, formed by transmembrane segments M1 and M2. In particular, the transmembrane densities (**Figure 4**) and the membrane-surface densities surround this calcium access funnel (asterisks in **Figure 5**). The cytoplasmic domain of PLN is highly basic and lies along the membrane surface (17). As such, it would be expected to perturb lipid packing adjacent to the calcium access funnel and transmembrane segments M1 and M2 of SERCA. This region of SERCA is highly mobile in the SERCA transport cycle, and the membrane perturbation may facilitate movement of transmembrane helices and thereby increase the turnover rate of SERCA. Thus, the interaction of PLN observed in the 2D crystals offers a plausible explanation for the effect on SERCA’s maximal activity.

As mentioned above, the affect of PLN on the maximal activity of SERCA is sensitive to PLN mutation. Merging our current findings with previous mutagenesis studies revealed that particular residues in the primary structure of PLN, when mutated to alanine, reduced the maximal activity of SERCA (31,44,46). The residues found to have the largest effect on the maximal activity of SERCA included Val^4^, Gln^5^, Asn^34^, Leu^37^, Met^50^, and Leu^51^. The first two residues, Val^4^ and Gln^5^, have minimal impact on PLN function (i.e. the ability to shift the apparent calcium affinity of SERCA), while the remaining residues are critical for PLN function (Asn^34^) and pentamer formation (Leu^37^, Met^50^, & Leu^51^). It is interesting that residues at the N-terminus (Val^4^ & Gln^5^) and C-terminus (Met^50^ & Leu^51^) of PLN are critical for this effect, suggesting that coordinated positioning of the cytoplasmic and transmembrane domains may play a role (Asn^34^ & Leu^37^ may also contribute). Residues were also found to increase the maximal activity of SERCA, when mutated to alanine, and these involved arginine residues in the cytoplasmic domain of PLN – Arg^9^, Arg^13^, and Arg^14^ (44). The modulation of charged residues in the cytoplasmic domain would be expected to alter the amphipathic membrane interactions of PLN. Together with the coordinated positioning of the cytoplasmic and transmembrane domains of PLN, these observations are consistent with the hypothesis that membrane perturbation may facilitate a more rapid turnover rate for SERCA.

## ACKNOWLEDGEMENTS

This work was supported by a grant from the Heart and Stroke Foundation (to H.S.Y.). J.P.G. was supported by a Canada Graduate Scholarship from the Canadian Institutes of Health Research and Alberta Innovates – Technology Futures. L.M.E.-F. was supported by a National Institutes of Health grant (R01GM120142).

## AUTHOR CONTRIBUTIONS

JPG and JOP performed the research, analyzed data, and contributed to the writing and editing of the manuscript; MJL contributed to the design and analysis of the data, and editing of the manuscript; LMEF performed and analyzed the protein-protein docking and molecular dynamics simulations; HSY designed the research, analyzed the data, and wrote the manuscript.

